# Endocannabinoids regulate mAChR-evoked theta rhythm IPSCs in hippocampus

**DOI:** 10.1101/022442

**Authors:** Ai-Hui Tang, Daniel A. Nagode, Bradley E. Alger

**Affiliations:** University of Maryland School of Medicine Department of Physiology

## Abstract

Exogenous cannabinoids can affect behaviorally relevant neuronal oscillations, but there is little evidence that endogenous cannabinoids (endocannabinoids, eCBs) can affect them, although it is unknown whether eCBs were generated during oscillations investigated in previous studies. In rat hippocampal slices, muscarinic receptor (mAChR) agonists stimulate the occurrence of persistent, rhythmic inhibitory post-synaptic currents (IPSC) activity and mobilize eCBs. We tested the hypothesis that mAChR-induced IPSCs would be modulated by concomitantly produced eCBs. With ionotropic glutamate receptors inhibited, mAChR agonist application triggered eCB-sensitive IPSCs that were enhanced in amplitude and frequency when a cannabinoid receptor antagonist was also present. There was also a highly significant increase in IPSC spectral power in the theta-frequency range. The data show that eCBs released by mAChRs modulate rhythmic IPSCs, and suggest that eCBs are candidate regulators of neuronal oscillations associated with eCB production *in vivo*.

**Significance statement:** Endocannabinoids (eCB) are ubiquitous and powerful modulators of neuronal activity, acting in the brain mainly via cannabinoid receptors (CB1Rs) on presynaptic nerve terminals, and inhibiting neurotransmitter release. Numerous neurotransmitters, including acetylcholine (ACh), stimulate eCB release. Behaviorally important neuronal oscillations can be suppressed by exogenous cannabinoids, suggesting that eCBs could regulate oscillations normally. Such regulation, if it occurred, would represent an important new dimension of eCB actions, but to date there is little evidence that eCBs do affect neuronal oscillations. Given the enrichment of eCBs and CB1Rs in brain regions whose behavioral roles depend on neuronal oscillations, including the hippocampus, it is important to understand whether, or how, the eCBs affect rhythms.

## INTRODUCTION

Neuronal electrical oscillations in the brain are associated with many behavioral actions and are considered essential for complex cognitive information processing. Exogenous cannabinoids are agonists at the major central cannabinoid receptor, CB1R, and disrupt both cognitive activity and neuronal oscillations (e.g., Robbe et al., 2006). Endogenous cannabinoids (endocannabinoids, eCBs) can suppress excitatory and inhibitory synaptic transmission by activating presynaptic CB1Rs (Kano et al., 2009; Castillo et al. 2012). Therefore eCBs should influence neuronal oscillations, yet to date there is little evidence that they do. eCBs do not affect the oscillations that have been investigated (e.g., Robbe et al. 2006; Mason and Cheer, 2009), but if the mechanisms of rhythm generation do not also stimulate the production and release of eCBs, there is no reason to expect eCBs to have an effect. Interneurons are key regulators of oscillations in the hippocampus, hence we tested the hypothesis that inhibitory oscillations in the rat hippocampal CA1 region that are generated by mAChRs, which also mobilize eCBs from pyramidal cells (Kim et al., 2002; Ohno-Shosaku et al., 2003) are regulated by eCBs.

Activation of mAChRs excites certain hippocampal interneurons (Cea-del Rio et al., 2011; Alger et al.,2014 for review) and CB1Rs are heavily expressed on nerve terminals of both the hippocampal perisomatic- and dendrite-targeting subclasses of cholescystokinin-expressing (CCK) interneurons; CB1Rs are essentially absent from other interneurons, including parvalbumin-expressing (PV) cells (Freund and Katona, 2007; Armstrong and Soltesz, 2012). The CB1R- and CCK-expressing (CB1R/CCK) interneurons are major sources of eCB-sensitive IPSCs (Lee et al., 2010).

Optogenetic stimulation of ACh release activates eCB-sensitive IPSCs in hippocampal CA1 (Nagode et al., 2011), and optogenetic silencing of glutamic-acid decarboxylase-expressing (Gad2) interneurons, which include the CB1R/CCK cells, reduced mAChR-induced, eCB-sensitive, theta-frequency IPSCs, whereas optogenetic silencing of PV interneurons did not affect them (Nagode et al. 2014). In contrast, optogenetic ACh release stimulates a subset of PV cells, which are are at least partially responsible, along with vasoactive intestinal peptide (VIP) cells, for the gamma-frequency IPSCs (Bell et al., 2015). CB1Rs are present at low levels on glutamatergic axons (Kano et al., 2009), but even when ionotropic glutamate receptors (iGluRs) are inhibited, mAChR agonists or optogenetic ACh release elicits bouts of theta frequency IPSPs/Cs in CA1 pyramidal cells (Alger et al., 2014, for review), demonstrating that these rhythms are independent of excitatory transmission. It appears that CCK cells are the primary source of mAChR-induced, theta-frequency, inhibitory oscillations.

mAChR-induced IPSCs are strongly suppressed by depolarization-induced suppression of inhibition (DSI, Pitler and Alger, 1992; Wilson and Nicoll, 2001; Fortin et al., 2004) which is mediated by eCBs (Wilson and Nicoll, 2001; Ohno-Shosaku et al., 2001). Therefore CB1R-expressing interneurons must be heavily involved in generating these IPSCs. It is possible that mAChR-induced IPSCs are sensitive only to eCBs mobilized during DSI, or alternatively, that these IPSCs are not fully suppressed by mAChR-generated eCBs. To distinguish between these hypotheses, we tested the prediction that mAChR-induced IPSC rhythms would be affected by a CB1R antagonist. If the IPSCs are only suppressed by eCBs released by DSI, then the antagonist should only affect DSI, but not other effects of the mAChR agonist on IPSCs, i.e., their amplitude or frequency. We elicited IPSCs with a pharmacological mAChR agonist (either carbachol or oxotremorine-M) in rat hippocampal slices and subsequently applied a CB1R antagonist (either AM251 or SR141716A). The results reveal a tonic influence of eCBs on rhythmic, theta-frequency mAChR-dependent IPSCs, suggesting that eCBs could contribute to regulation of certain *in vivo* oscillations.

## METHODS

All animal experiments were approved by the [Author University] Institutional Animal Care and Use Committee and were conducted in accordance with national and international guidelines to reduce pain and suffering as well as the numbers of animals used. We used tissue from 5-6 week old male Sprague-Dawley rats (Charles River, http://www.criver.com/products-services/basic-research/find-a-model/sprague-dawley-rat). Animals were deeply sedated with isoflurane and decapitated. Slices, 400 μm thick, were cut on a Vibratome (VT1200s, Leica Microsystems) in an ice-cold extracellular recording solution (ACSF) containing (mM): 130 NaCl, 3 KCl, 2.5 CaCl_2_, 2 MgSO_4_, 1 NaH_2_PO_4_, 25 NaHCO_3_, and 10 glucose, and was bubbled with 95% O_2_, 5% CO_2_, pH 7.4. Slices were stored in a holding chamber on filter paper at the interface of ACSF and a moist, oxygenated atmosphere at room temperature (22°C) for >1 h before transfer to the recording chamber (RC-27L, Warner Instruments).

Whole-cell recordings were made with a Nikon E600 microscope. CA1 pyramidal cells were voltage-clamped at -70 mV with 3-5 MΩ glass pipette filled with (mM): 90 CsCH_3_SO_4_, 1 MgCl_2_, 50 CsCl, 2 MgATP, 0.2 Cs_4_-BAPTA, 10 HEPES, 0.3 Tris GTP, and 5 QX314, 280–290 mOsm, pH 7.2. Series resistances were monitored and recordings with a change of >20% were discarded. Data were collected with Axopatch 200B amplifiers (Molecular Devices), filtered at 2 kHz, and digitized at 5 kHz with Digidata 1440 (Molecular Devices) and Clampex 10 software (Molecular Devices).

NBQX (10 μM) and D-AP5 (20 μM) were present to block ionotropic glutamate receptors. To induce spontaneous inhibitory postsynaptic currents (sIPSC), we applied the selective mAChR agonist, oxotremorine-M (Oxo-M, 2 μM) or the non-selective cholinergic agonist, carbachol (CCh, 5 μM). CB1R activation was prevented by bath application of one of the inverse agonists, SR141617A (rimonabant, 2 μM) or AM251 (5 μM). AM251 and SR141617A were dissolved in ethanol (5-10 mM stock). The amplitudes and inter-IPSC intervals were measured with MiniAnalysis (Synaptosoft) using a detection threshold of 8 pA. The power analysis was performed in Clampfit 10. Statistical tests were done with Sigmaplot 7.0. Cumulative distributions were compared with the Kolmogorov–Smirnov (K–S) test (http://www.physics.csbsju.edu/stats/KS-test.n.plot_form.html). Power spectra between groups were compared with two-way ANOVA. Averaged power amplitudes were tested using paired t-test. The significance level for all tests was p < 0.05 (∗). Effect size was calculated with Excel.

## RESULTS

Whole-cell recordings from pyramidal cells in CA1 hippocampal slices in which iGluRs were pharmacologically blocked, confirmed that an mAChR agonist induces persistent IPSC activity that lasts for tens of minutes (Fig. 1A). Application of a 2-s voltage step from the holding potential of -70 mV to 0 mV caused major but transient suppression of these IPSCs (i.e. DSI) as previously reported (cf., Pitler and Alger, 1992; Wilson and Nicoll, 2001; Fortin et al., 2004; Fig. 1A). DSI could be elicited repeatedly without significant change for the duration of a recording, demonstrating that the mAChR-induced IPSCs can be blocked by eCBs produced by DSI, and therefore that mAChR-stimulated, CB1R-expressing interneurons remain capable of releasing GABA. The question is whether or not mAChR-generated eCBs have any effect on the ongoing IPSCs, as the hypothesis predicts.

To test this, we first induced the persistent occurrence of large, rhythmic IPSCs in CA1 pyramidal cells with bath-application of the selective mAChR-agonist, Oxo-M, and then applied the CB1R inverse agonist/antagonist, SR141716A. DSI was induced at 2-min intervals throughout. Once the IPSCs and DSI had been stable for ~15 min, we applied SR141716A, at 2 μM, via the perfusion solution and left it in for the remainder of the recording together with Oxo-M. We observed that SR141716A gradually reduced the IPSC suppression by DSI, confirming previous results on evoked IPSCs (Wilson and Nicoll, 2001; Kim et al., 2002). However, concomitant with the disappearance of DSI, the IPSC amplitudes gradually increased as SR141716A took effect (e.g., Fig. 1A). Fig. 1B illustrates this experiment in more detail, with every dot representing a single IPSC amplitude. The entire population of IPSCs did not simply increase in amplitude, but a subgroup of the IPSCs occurring during the maximal SR141716A block of DSI is much larger than those before its application. Examination of portions of this trace on an expanded time scale indicates that essentially all of the largest IPSCs were abolished by DSI in the control period, but are unaffected by the DSI-stimulation after SR141716A took effect. Figure 1C illustrates the time course of the DSI reversal in all 7 cells from 5 animals in this group. Group data of all IPSCs (n =700) from 7 cells (same ones as in Fig. 1C) treated with Oxo-M followed by Oxo-M plus SR141716A is shown in Fig. 1D1 (p < 0.001, K-S test). Data (n = 600 IPSCs) from a separate group of 6 cells from 5 animals treated with the non-specific mAChR agonist, carbachol (CCh, 5 μM) and a different CB1R antagonist, AM251 (2 μM) is shown in Fig. 1E1 (p < 0.001, K-S test). By revealing that the CB1R antagonists cause a rightward shift in the cumulative frequency plots of IPSC amplitudes, these results are consistent with the prediction that stimulation of mAChRs mobilizes eCBs and exerts a partial, tonic suppressive effect on the IPSC amplitudes.

**Figure 1.**
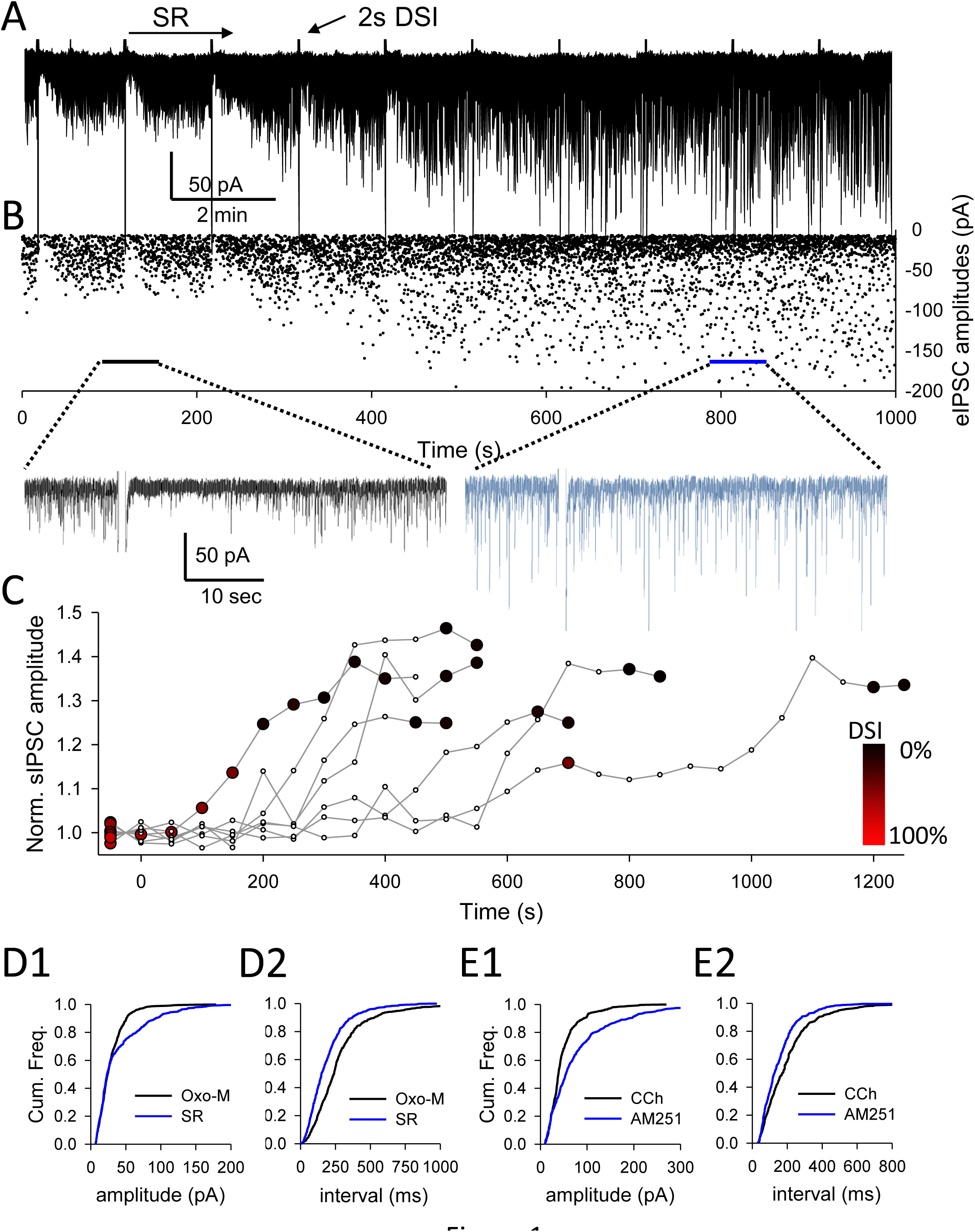
CB1R antagonism uncovers a tonic, CB1R-dependent partial suppression of IPSC activity induced by mAChR activation. A) Whole cell recording from a CA1 pyramidal cell shows the persistent barrage of IPSCs (rapid downward deflections) induced by Oxo-M, with the IPSCs being markedly reduced every 2-min (DSI) induced by a 2-sec voltage step to 0 mV (V_h_ = -70 mV). iGluR antagonists were in the bathing solution in all experiments. The CB1R antagonist, SR141716A (2 μM), was added to the bath perfusion at the time indicated by the arrow and kept in for the remainder of the experiment. As SR141716A reduced DSI, it increased the IPSC amplitudes. Indicated portions of the traces are shown on an expanded time scale below. B) Time course of the amplitudes of all sIPSCs for experiment shown in A), each sIPSC amplitude represented by a dot. C) Time course of DSI reversal during wash-in of SR141716A for all 7 cells in this group. IPSC amplitudes normalized to peak depression during DSI, and increase above that level as SR141716A washes in. Maximal DSI (100%) occurs at peak IPSC depression, color coded in red. D) Cumulative frequency plots of group data (n=7 cells, same cells as in 1C) 700 IPSCs total) showing the distribution of amplitudes (D1) and inter-event intervals (D2) before (black traces) and after (blue traces) addition of SR141716A to slices in which Oxo-M had induced persistent IPSC activity. The rightward shift in B1 and leftward shift in B2 indicates that SR141716A increased the amplitudes and frequency of the IPSCs, respectively. E) Cumulative frequency plots of group data (n = 6 cells, different cells than in B, 600 IPSCs) that were treated with CCh and AM251. E1) and E2) analogous to D1 and D2. Results from the two groups of cells are essentially identical.

The mAChR-generated eCBs might influence IPSC frequency as well, so to check for this we measured the IPSC inter-event interval distributions. Figure 1D2 shows that SR141716A causes a leftward shift in the cumulative frequency plot of IPSC inter-event intervals; i.e., in SR141716A, IPSCs occurred at shorter intervals, which represents a higher ongoing frequency (p < 0.001, K-S test). Figure 1E2 shows that AM251 had a similar effect on CCh-induced IPSCs (p < 0.001, K-S test). Thus the data suggest that persistent mobilization of eCBs by pharmacological application of mAChR agonists has two effects: it reduces IPSC amplitudes and slows their frequency; both effects are revealed by a CB1R antagonist. The implications of these results are considered further in the Discussion.

Because the mAChR-mobilized eCBs suppress the frequency of IPSCs, they could also affect the IPSC rhythmicity. We tested this with power spectral analyses of mAChR-mediated IPSCs in the absence and presence of a CB1R antagonist (Fig. 2A). The graph in Fig. 2A shows that Oxo-M predominantly triggered the occurrence of IPSCs with a peak power at just under 3Hz. After SR141716A was added to the bathing solution, the spectral power increased greatly throughout the range of 2-15Hz, and the peak power increased to ∼4 Hz. Fig. 2B illustrates that both SR141716A (left plot, p = 0.008, paired t-test) and AM251 (right plot, p = 0.019, paired t-test) caused highly significant increases in the total spectral power over the range of 2-15Hz. Fig. 2C illustrates that the CB1R antagonists increased the peak frequency of the IPSCs from the same cells.

**Figure 2.**
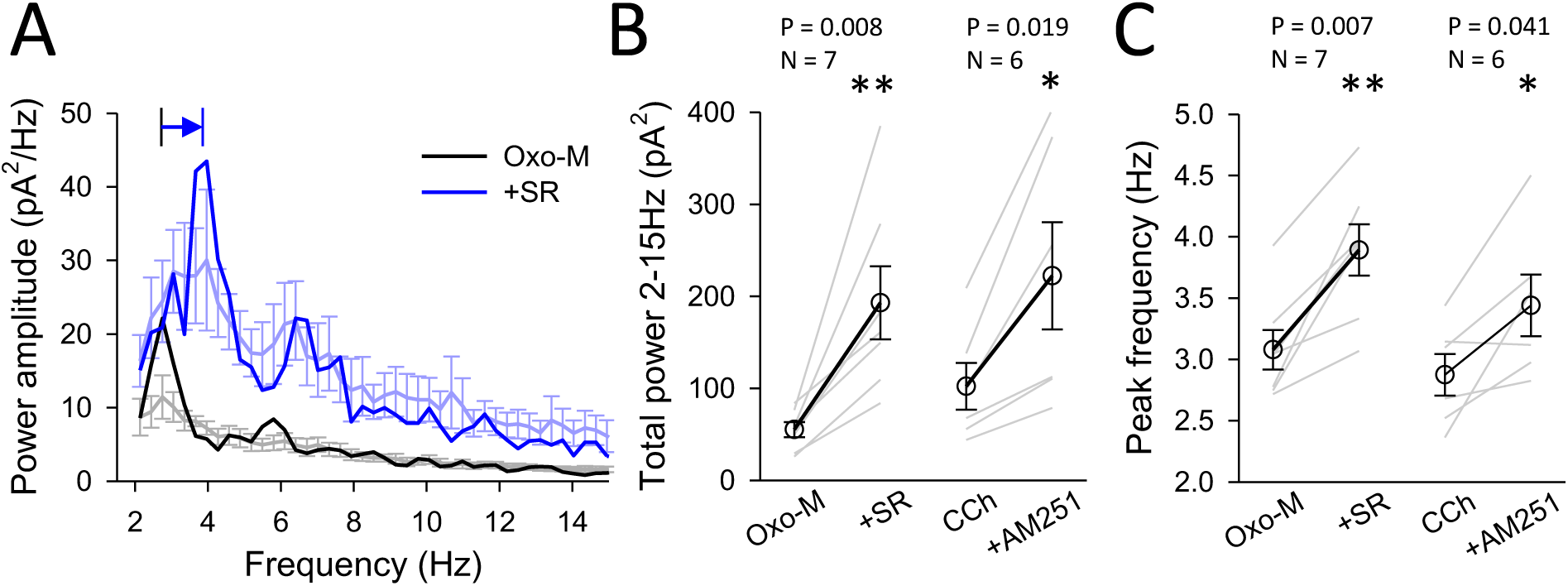
CB1R antagonism increases spectral power in the theta frequency range. A) Group data showing power spectral analysis for 7 cells from rat slices treated with Oxo-M before and after application of SR141716A. Light traces with error bars were pooled results. Thick traces represent the data from the example traces in Figure 1. B) Shows total power summated over the theta range from individual slices treated with Oxo-M and then Oxo-M plus SR141716A (left) or CCh and CCh plus AM251 (right). C) Shows peak power measured in the same groups as in 2B.

## DISCUSSION

Despite copious evidence that mAChRs can both stimulate vigorous firing of certain interneurons and mobilize eCBs, it has been unclear whether or not mAChR-induced IPSCs are themselves affected by mAChR-mobilized eCBs (Robbe et al., 2006; Mason and Cheer, 2009; Alger et al., 2014). Because these IPSCs are readily inhibited by the eCB-mediated process, DSI, they must come from CB1R-expressing interneurons. However, a lingering possibility was that the mAChR-induced IPSCs might only be susceptible to the eCBs that are produced during DSI, and not those produced by mAChR stimulation. While 2-arachidonyl-glycerol (2-AG) is probably the eCB that is responsible for both DSI and mAChR effects, the biochemical pathways for 2-AG mobilization induced by DSI and by mAChRs are qualitatively different (i.e., PLCβ is part of the mAChR pathway, but not the DSI pathway; Hashimotodani et al., 2005), and it was conceivable that their physiological effects were also different. We reasoned that if mAChR-induced eCBs reduced, but did not abolish, the mAChR-induced IPSCs, then a CB1R antagonist would alter the IPSC amplitudes, their frequency, or both, in addition to blocking DSI. Our results show that CB1R antagonists consistently enhanced IPSC frequency, suggesting that concomitantly with triggering IPSC activity, mAChRs partially suppress this activity by releasing eCBs.

The CB1R antagonists appear to have no effect on small IPSCs: the cumulative frequency plots of IPSC amplitudes before and after CB1R antagonist application essentially overlap for IPSCs below ∼25 pA (Figs. 1B1, 1C1). The CB1R antagonists did increase the proportion of larger amplitude IPSCs, which would be consistent with the suggestion that it is mainly the perisomatic IPSCs that are affected by eCBs. We cannot distinguish between the possibilities that the CB1R-antagonists increased the quantity of GABA released from each terminal, an analog mechanism, or that they increased the number of terminals releasing GABA by “unsilencing” some of them (Losonczy et al., 2004; Neu et al., 2007), a digital mechanism. Either explanation could account for the increase in amplitude and decrease in interval of the large IPSCs (Figs. 1D, 1E), and will have to be distinguished in future work.

At first glance the idea of a system having two opposing effects on IPSCs might seem self-defeating, but in fact interplay of opposing influences is common in control systems. Temperature stabilization in modern buildings is often carried out by playing cooling and heating functions off each other. This dual ability enables the central controller to exert maximal effects in either direction rapidly and maintain the setpoint efficiently. The mAChR system stimulates firing in CB1R+ interneurons (among others) and releases eCBs which partially suppresses their GABA release. But the strength of eCB-mediated inhibition is modulated by the firing frequency of the interneuron (Losonczy et al., 2004; Földy et al., 2006; Karson et al., 2009) -- the faster the firing, the weaker the inhibition. The net rate at any moment will reflect a balance of the two forces. The dual effects of the mAChRs could provide a highly sensitive way of regulating the degree of inhibition exerted by the interneurons on their target cells. Of course, within *in vivo* neuronal circuits there are other excitatory and inhibitory influences that will also alter the firing of the interneuron – either up or down – and this in turn will alter the degree to which the eCBs can inhibit its GABA release.

Our experiments were done on a simplified model of neuronal oscillations and it is unknown to what extent these results are relevant to behaviorally relevant oscillations in vivo. There is good evidence that certain forms of *in vivo* oscillations are driven by GABAergic inhibition and are atropine-sensitive, implying that mAChRs play a major role in generating them; other forms of theta rhythm are not atropine-sensitive (Buzsaki, 2002). Undoubtedly only certain forms of oscillations will be sensitive to modulation by eCBs. Indeed, theta, gamma and ripple oscillations generated *in vivo* in head-restrained and freely moving animals (Robbe et al., 2006) or epileptifom discharges induced by kainic acid in urethane-anesthetized animals (Mason and Cheer, 2009) are not altered by administration of a CB1R antagonist, suggesting the eCBs do not modulate them. Note that kainic acid inhibits GABA release (Fisher and Alger, 1984; Carta et al., 2014) including GABA release from CB1R-expressing interneurons (Lourenco et al., 2010), which could account for the insensitivity of kainic-acid induced oscillations to CB1R antagonists. Identified interneurons active during kainic acid induced gamma rhythms in CA1 are not perisomatic targeting cells and do not express CB1Rs (Craig and McBain, 2015). We predict that *in vivo* oscillations that depend on CB1R+ interneurons and are induced by mechanisms that mobilize eCBs, such as atropine-sensitive theta, will be influenced by eCBs.

